# Genetic and phenotypic diversity of wine-associated *Hanseniaspora* species

**DOI:** 10.1101/2025.04.09.648064

**Authors:** Cristobal A. Onetto, Chris Ward, Cristian Varela, Laura Hale, Simon A. Schmidt, Anthony, Borneman

## Abstract

The genus *Hanseniaspora* includes apiculate yeasts commonly found in fruit- and fermentation-associated environments. Their genetic diversity and evolutionary adaptations remain largely unexplored despite their ecological and enological significance. This study investigated the phylogenetic relationships, genome structure, selection patterns, and phenotypic diversity of *Hanseniaspora* species isolated from wine environments, focusing on *Hanseniaspora uvarum*, the most abundant non-*Saccharomyces* yeast in wine fermentation. A total of 151 isolates were sequenced, including long-read genomes for representatives of the main phylogenetic clades. Comparative genomics revealed ancestral chromosomal rearrangements between the slow- (SEL) and fast-evolving (FEL) lineages that could have contributed to their evolutionary split, as well as significant loss of genes associated with mRNA splicing, chromatid segregation and signal recognition particle protein targeting specifically in the FEL lineage. Pangenome analysis within *H. uvarum* identified extensive copy number variation (CNV), particularly in genes related to xenobiotic tolerance, nutrient transport and metabolism. Investigation into the selective landscape following the FEL/SEL divergence identified diversifying selection in 229 genes in the *Hanseniaspora* FEL lineage, with significant enrichment in genes within the lysine biosynthetic pathway, suggesting a key role for this amino acid in early FEL adaptation. In *H. uvarum*, signatures of recent positive selection were detected in genes linked to sulphur assimilation, sterol biosynthesis and glycerol production, indicating potential adaptation to the stresses imposed by grape and wine fermentation. Furthermore, phenotypic screening of 113 isolates revealed substantial intraspecific diversity, with specific species exhibiting enhanced ethanol, osmotic, copper, SO₂, and cold tolerance. These findings provide novel insights into the genomic evolution and functional diversity of *Hanseniaspora*, expanding our understanding of yeast adaptation to wine fermentation and laying the foundation for targeted gene investigations within this important genus.

## 1. Introduction

The genus *Hanseniaspora* consists of a group of apiculate yeasts commonly found in natural environments, including fruit (Chanprasartsuk et al. 2010, Prada and Pagnocca 1997, Ramirez-Castrillon et al. 2019, Vadkertiová et al. 2012, Vegas et al. 2020), flower/nectar (Čadež et al. 2014, Cadez and Smith 2011, Klaps et al. 2020) and fermentation ecosystems such as wine (Chanprasartsuk et al. 2010, Lorenzini et al. 2018, Onetto et al. 2024). These yeasts are characterised by a preference for sugar-rich environments, bipolar budding and reduced genome sizes compared to other members of the Saccharomycetales. Among the most studied species in winemaking are *Hanseniaspora uvarum*, *Hanseniaspora guilliermondii*, and *Hanseniaspora vineae*, which have been isolated from the surfaces of grape berries and the early stages of wine fermentation (Steenwyk et al. 2019, van Wyk et al. 2024).

*Hanseniaspora* species, particularly *H. uvarum*, play a significant role in the early stages of spontaneous wine fermentation, where they influence both the microbial succession and chemical composition of wine (Ciani et al. 2006, Onetto et al. 2025, Onetto et al. 2024, Pourcelot et al. 2025, Snyder et al. 2024). While traditionally regarded as spoilage organisms due to the propensity of some strains to produce high levels of acetic acid and ethyl acetate (Fleet 1990, Fleet et al. 1984), recent studies have highlighted potential positive contributions to wine aroma complexity (Mestre et al. 2019, Rossouw and Bauer 2016, Santos et al. 2016). Certain *Hanseniaspora* species, particularly *H. vineae*, have been shown to enhance the production of desirable volatile compounds such as esters and higher alcohols, which contribute to fruity and floral aromas (Carrau et al. 2023, Varela 2016). As a result, there is increasing interest in the potential application of *Hanseniaspora* species to enhance wine aroma in controlled mixed fermentations (van Wyk et al. 2024).

*Hanseniaspora* species can be broadly characterised into two main phylogenetic lineages with distinct evolutionary rates: a slow-evolving lineage (SEL) and a fast-evolving lineage (FEL). Both lineages display extensive gene loss, particularly in pathways related to DNA repair and cell-cycle regulation (Steenwyk et al. 2019). A recent study by Haase et al. (2024) revealed that *H. uvarum* (FEL) has lost core histone genes typically conserved in Saccharomycetales, while acquiring novel cis-regulatory elements that may have compensated for these losses. These adaptations have implications for yeast cell-cycle progression and competitiveness within the fermentation ecosystem (Lax and Gore 2023, Onetto et al. 2025, Schwarz et al. 2022).

Despite their ecological and oenological significance, the genetic diversity and evolutionary dynamics of *Hanseniaspora* species remain underexplored. Population-level studies have identified some degree of spatial and temporal genetic clustering in *H. uvarum* isolates from winemaking environments (Albertin et al. 2016, Saubin et al. 2020). However, gene content variation, genetic diversity, and selection patterns remain largely unknown due to the absence of whole-genome sequencing studies across multiple isolates of this genus.

Understanding the genetic diversity of *Hanseniaspora* species is crucial for fundamental microbiology and applied oenology. This study addresses these knowledge gaps by characterising the genetic and phenotypic diversity of *Hanseniaspora* populations sourced from winemaking environments and provides insights into the gene content variation, genome synteny, selection patterns and phenotypic diversity of this ecologically and oenologically significant genus.

## 2. Material and Methods

### 2.1 *Hanseniaspora* isolates

A total of 151 *Hanseniaspora* isolates were obtained from the AWRI culture collection and selected based on prior species identification via internal transcribed spacer (ITS) sequencing. Most of the isolates were obtained from spontaneous grape juice fermentations across Australia as part of a large bioprospecting initiative (Onetto et al. 2024). Additional isolates from diverse regions worldwide were also included in this study (Table S1).

### 2.2 DNA extraction and genome sequencing

For short-read sequencing, yeast DNA was isolated by lysis of protoplasts formed through zymolyase digestion and potassium acetate precipitation, as previously described (Davis et al. 1980). Library preparation and sequencing were performed in the Ramaciotti Centre for Genomics (University of New South Wales, Sydney, Australia). Sequencing libraries were prepared using the Nextera DNA flex kit and sequenced with an Illumina NovaSeq 6000 using 2 x 150 bp chemistry on an SP flow cell.

For long-read sequencing, DNA was extracted using a Gentra Puregene Yeast/Bact DNA extraction kit (Qiagen, Australia). Sequencing libraries were prepared using the SQK-NBD114-24 kit and sequenced in FLO-PRO114M flow cell (Oxford Nanopore Technologies, Oxford, UK). Pod5 files were basecalled using Dorado (https://github.com/nanoporetech/dorado) with the Sup model v4.3.0.

### 2.3 Genome assembly, annotation and synteny

Short-reads were quality trimmed using fastp (Chen et al. 2018) and genome assemblies were performed using MEGAHIT (Li et al. 2015). Long-read genome assemblies were performed using Flye (Kolmogorov et al. 2019) and polished using Medaka (https://github.com/nanoporetech/medaka).

Gene prediction of genome assemblies was performed following the funannotate pipeline (Palmer and Stajich 2017), including Genemark-ES v. 4.68 (Ter-Hovhannisyan et al. 2008), SNAP (Korf 2004), Augustus (Stanke and Waack 2003) and Glimmerhmm (Delcher et al. 1999) annotations trained using BUSCO (Manni et al. 2021). Functional annotations were performed using the UniProt database (2021_02), Interproscan 5 (Jones et al. 2014), Pfam (Mistry et al. 2020), and dbCAN (Zheng et al. 2023) databases.

Protein orthology based synteny between the long-read genome assemblies was performed using GENESPACE (Lovell et al. 2022).

Publicly available hybrid *H. pseudoguilliermondii* x *H. opuntiae* short reads (Saubin et al. 2020) and hybrid strains from this study were mapped to the long read reference *H. pseudoguilliermondii* x *H. opuntiae* (wild-127) using bwa mem (Li 2013). Read depth was then calculated using samtools (Danecek et al. 2021). Significant changes in local coverage were determined using the difference between the local vs genome-wide read depth distribution (Ward et al. 2024).

Structural variants were determined by mapping *H. pseudoguilliermondii* x *H. opuntiae* wild-45 using nglmr (Sedlazeck et al. 2018) and calling with sniffles (Sedlazeck et al. 2018). Structural variants were then manually investigated in IGV (Robinson et al. 2011) to identify well-supported variants.

### 2.4 Phylogenetic analysis, variant calling, ploidy and copy number variation

Orthogroups were identified using Orthofinder (Emms and Kelly 2018b) and aligned using MUSCLE (Edgar 2004). Translation-informed codon alignments were obtained using pal2nal (Suyama et al. 2006) and trees were generated for each orthogroup using IQ-TREE (Nguyen et al. 2014) with the best-fit model for each tree automatically selected using ModelFinder (Kalyaanamoorthy et al. 2017). The species tree was inferred using all orthogroups with STAG (Emms and Kelly 2018a).

For variant calling, short-read sequences were mapped to their corresponding long-read reference assembly using BWA (Li 2013) and duplicates were removed using picard (Toolkit 2019). Variant calling was performed using Bcftools (Danecek et al. 2021). Ploidy was estimated using k-mers with Smudgeplot (Ranallo-Benavidez et al. 2020) and SNPs as described previously in Ward et al. (2024).

Copy number variants (CNV) were identified using the python package CNVpytor (Suvakov et al. 2021) with telomere and centromeres regions masked before read depth estimation.

### 2.5 Pangenome graph construction and gene graph-based presence-absence variations

Pangenome graph construction for *H. uvarum* was performed using the Minigraph-Cactus pipeline (Hickey et al. 2024) using the chromosome-level genome assemblies of isolates wild-6, wild-104 and wild-115. After graph construction, short-read sequences of the remaining *H. uvarum* isolates were mapped to the graph using vg-giraffe (Sirén et al. 2021) and bam files extracted for each sample and reference using vg-surject (Garrison et al. 2018). The coverage for each ORF for each sample-reference combination was calculated with BEDtools (Quinlan and Hall 2010) and ORFs with less or higher than 0.8 coverage were considered dispensable or core, respectively. All dispensable and core genes in all sample-reference combinations were deduplicated based on protein homology using a combination of BLASTp (Camacho et al. 2009) and OrthoMCL (Li et al. 2003).

### 2.6 Selection testing

Screens for selective sweeps within the *H. uvarum* population were performed using the μ statistic as previously described by (Onetto et al. 2022) ultilising the multi-sample vcfs obtained from section 2.4 after masking for telomeric, centromeric and repetitive regions. The μ statistic relies on multiple signatures of a selective sweep, including expected reduction of variation in the region of a sweep, shifts in site frequency spectrum (SFS) and emergence of localized LD patterns on each side of the beneficial mutation (Alachiotis and Pavlidis 2018). A percentile threshold of 99.9% was applied to the μ statistic, and top 0.1% scored overlapping windows were merged to detect sweep regions.

For episodic diversifying selection, single copy orthogroups between the *H. uvarum, H. opuntiae, H. guilliermondii, H. valbyensis, H. vineae* and *H. osmophila* reference genomes, identified using Orthofinder (Emms and Kelly 2018b), were codon aligned using TranslatorX (Abascal et al. 2010). Stop codons were then removed using TrimAl (Capella-Gutiérrez et al. 2009) and codon alignments were tested for diversifying selection using the Busted-PH model from HyPhy (Murrell et al. 2015, Pond et al. 2004). A binary tree with the FEL branch set as the foreground and the SEL branch as the background was utilised for all trees. The output of Busted-PH was then parsed to identify orthogroups with significant (p < 0.05) evidence for diversifying selection specific to the FEL branch, specific to the SEL branch and along both the FEL and SEL branches but with significant differences in ω distributions.

### 2.7 Phenotype screening

Phenotype screening was performed in duplicate, as previously described in (Varela et al. 2023). Briefly, colonies were pinned onto synthetic complete (SC) agar medium (20 g/L of glucose, 1.7 g/L yeast nitrogen base without ammonium sulfate or amino acids, 5 g/L ammonium sulfate and 20 g/L agar) and grown at 28 °C for 5 days. These plates were used as source plates for screening colonies under different environmental conditions, including SC agar media containing 3%, 6% or 12% (v/v) ethanol, 20% or 40% (w/v) glucose, 0.2 mM copper sulfate and 1 mM sodium sulfite. Hydrogen sulfide (H_2_S) production was assessed in SC agar media containing copper, since H_2_S reacts with copper generating CuS, which turns colonies brown. Agar plates were incubated at 28 °C for 5 days and then photographed using the PhenoBooth (Singer Instruments, Somerset, UK). Photos were used to quantify colony sizes and measure colony colour with the software PhenoSuite v2.20.504.1 (Singer Instruments, Somerset, UK).

The sensitivity of isolates to environmental conditions was quantified by expressing the ratio of the colony size for each test condition over the colony size on SC agar medium. For sulfite tolerance, the ratio was obtained from the colony size in media with 4.5 g/L tartaric acid and 1mM sodium sulfite over the colony size on SC media containing only 4.5 g/L of tartaric acid. H_2_S production was expressed as the ratio of the colony brightness in SC media over the colony brightness on media containing copper. The phenotype data for all the strains is available in Table S2.

Genome-wide association analysis (GWAS) utilising the phenotype data of the sequenced *H. uvarum* strains and the multi-sample vcf produced in section 2.4 was performed with GEMMA (Zhou and Stephens 2012), utilising a likelihood ratio test (option -lmm 2) and accounting for population structure using the centred relatedness matrix (option -gk 1).

### 2.8 Data availability

The sequencing data and genome assemblies included in this study are publicly available in the NCBI repository under BioProject PRJNA1231130.

## 3. Results and Discussion

### 3.1 Phylogenetic and genetic diversity of *Hanseniaspora* isolates

Whole-genome sequencing was performed on a set of 150 *Hanseniaspora* isolates that included representatives of the previously described SEL and FEL lineages (Steenwyk et al. 2019). These genomes, in addition to 21 publicly available reference genomes (Čadež et al. 2019, Čadež et al. 2021, Giorello et al. 2014, Langenberg et al. 2017, Opulente et al. 2024, Riley et al. 2016, Seixas et al. 2018, Shen et al. 2018, Sternes et al. 2016) were used to define a set of 1353 orthogroups that were subsequently utilised for gene-tree based phylogenetic reconstruction (Figure 1). The resulting structure was consistent with previously published *Hanseniaspora* phylogenies, with the FEL and SEL lineages clearly defined (Steenwyk et al. 2019). Phylogenetic placement indicated that the sequenced isolates corresponded to *H. uvarum*, *H. opuntiae, H. guilliermondii*, *H. valbyensis*, *H. vineae*, *H. osmophila*, and *H. occidentalis* (Figure 1).

**Figure 1.**
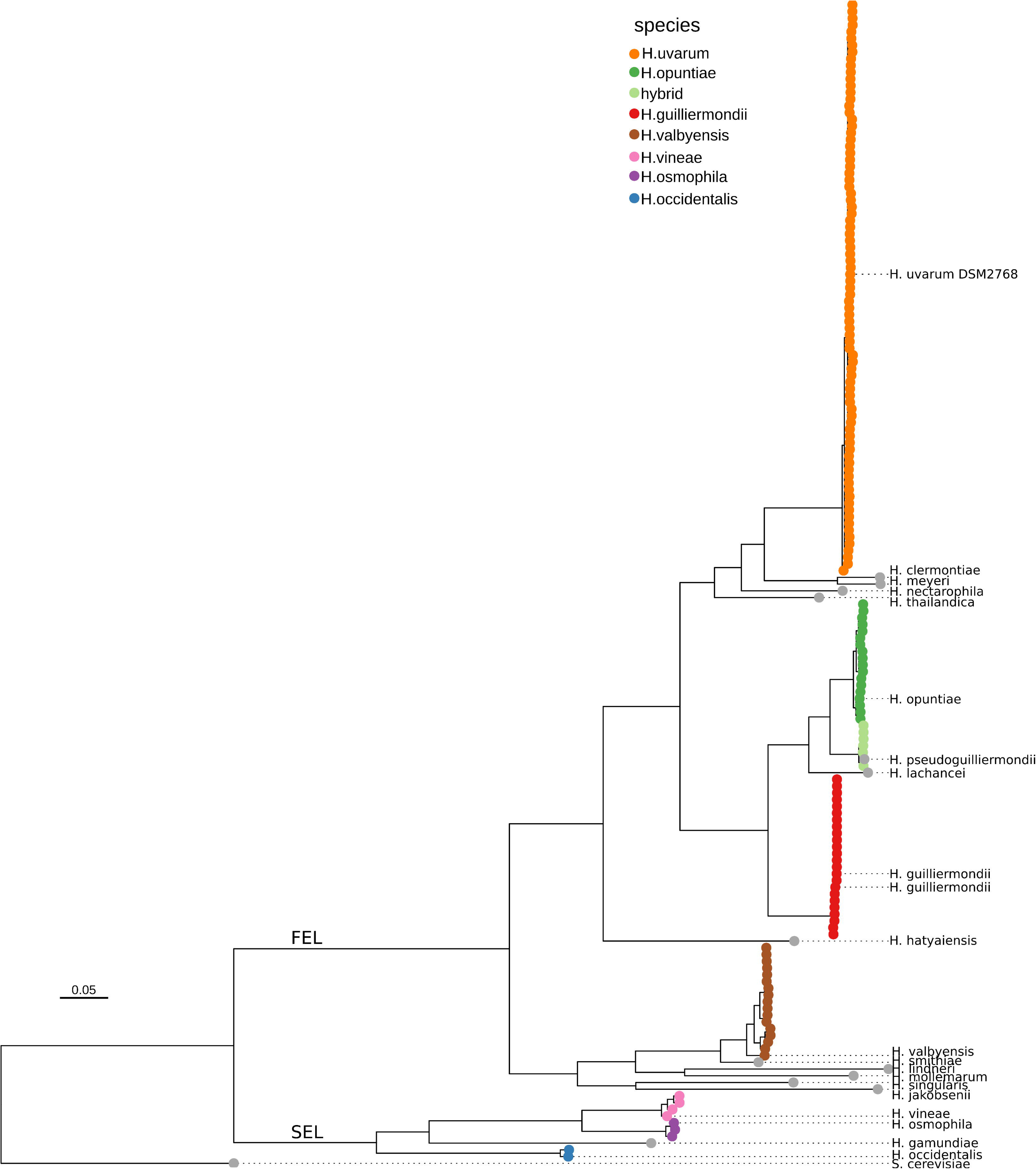
Phylogenetic tree inferred from 1353 codon-aligned orthologue gene trees with all species present using the STAG algorithm. Labelled strains correspond to genomes available in NCBI and unlabelled strains were sequenced for this study. Branch lengths represent substitutions per site.

Consistent with previous findings (Steenwyk et al. 2019), species within the SEL showed significantly larger genome assemblies than those in the FEL (Figure S1a), with an average 1n genome assembly size of 11.51 ± 0.15 Mb for SEL species compared to 9.06 ± 0.47 Mb for FEL species.

To precisely estimate genetic diversity, ploidy, and chromosomal synteny among *Hanseniaspora* species, long-read genome assemblies were generated for 19 representative strains from each clade. Analysis of the long-read assemblies for isolates initially classified as *H. pseudoguilliermondii* revealed unusually large genome assemblies and further investigation determined these isolates to be interspecific hybrids of *H. opuntiae* and *H. pseudoguilliermondii* (Figure S2). This hybridization event has previously been reported in strains isolated from France, Slovakia, Algeria, and Mayotte Island (Saubin et al. 2020), suggesting it may be a recurrent phenomenon.

The long-read assembly of one hybrid strain that contained near-complete *H. opuntiae* and *H. pseudoguilliermondii* phased haplotypes (Figure S2) was used to generate read-depth plots for hybrid assessment. All isolates from this study were predicted to be allotriploids, derived from an ancestral *H. opuntiae* (2n) × *H. pseudoguilliermondii* (1n) genome (Figure S3, Table S4). These ploidy observations are consistent with previous findings for other hybrid isolates (CLIB 3101, DBVPG 5828, and CCY46-1-3, Figure S4) (Saubin et al. 2020). One isolate (wild-45) exhibited multiple structural variations between parental haplotypes (Figure 2a), while another (wild-44) showed a near-complete chromosomal loss of the *H. opuntiae* haplotype, which was counterbalanced by duplications in the *H. pseudoguilliermondii* haplotype (Figure S3d, contig_55-contig_56). Mapping of previously published short-read data from hybrid strains (Saubin et al. 2020) also revealed similar read-depth changes indicative of structural rearrangements (Figure S4), suggesting that chromosomal rearrangements between haplotypes are a common feature of these hybrids.

**Figure 2.**
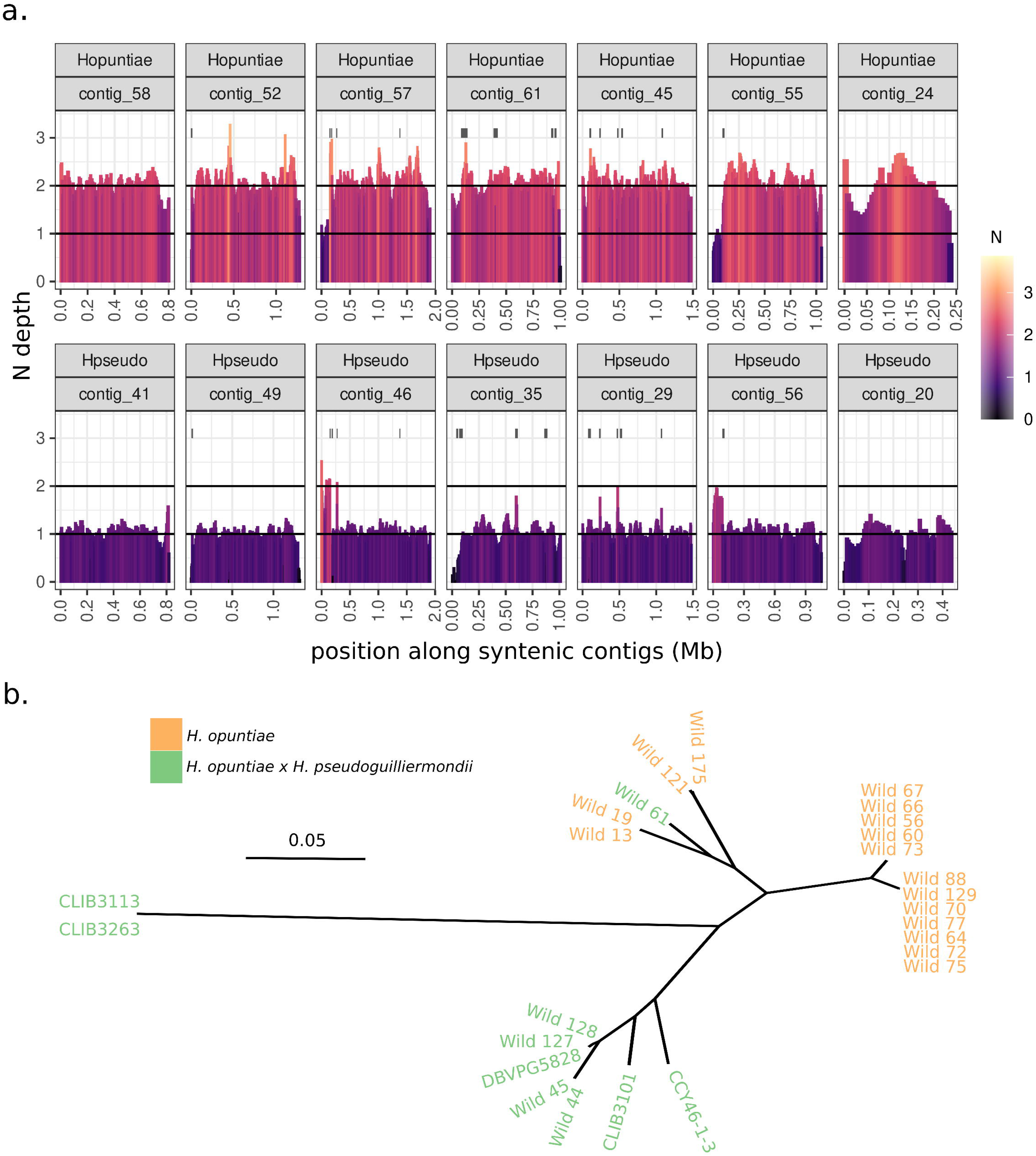
Investigating structural variation and evolutionary history of H. opuntiae x H. pseudoguilliermondii hybrids. **a.** Relative N read depth of hybrid wild-45 strain long-reads mapped to the wild-127 H. opuntiae x H. pseudoguilliermondii reference genome. Syntenic contigs of H. opuntiae and H. pseudoguilliermondii origin have been separated to show ploidy of the H. opuntiae and H. pseudoguilliermondii genome haplotypes within the wild-45 long reads. Black lines above the read depth histogram indicate the predicted positions of translocations between H. opuntiae and H. pseudoguilliermondii genome haplotypes, predicted using long reads. **b.** Maximum likelihood phylogenetic reconstruction of variants called against wild-127 H. opuntiae contigs. H. opuntiae and hybrids generated in this study are compared against publicly available H. opuntiae x H. pseudoguilliermondii hybrids.

Mating events leading to interspecific hybrids are prevalent in *Saccharomycotina* yeast species, including industrially relevant hybrids such as the brewing yeast *Saccharomyces pastorianus* and *Saccharomyces bayanus*, where they are believed to provide strong selective advantage in novel, anthropomorphic environments (Gabaldón 2020, Gallone et al. 2019, Gardner et al. 2024, Langdon et al. 2019). While genome sequence of the type strain isolated from orange juice has been shown to represent a “pure” strain of *H. pseudoguilliermondii* (Shen et al., 2018), all wine/grape isolates originally believed to be *H. pseudoguilliermondii* have been identified in this work to be interspecific hybrids of *H. opuntiae* and *H. pseudoguilliermondii*. This suggests that this hybridisation event represents an adaptation to the grape environment. Accordingly, phenotyping of these hybrids under common wine-related stress conditions revealed that they exhibited fitness levels comparable to, or better than, many *Hanseniaspora* species, including isolates of *H. opunti*ae (see later, Figure 5).

Allotriploid hybrids, while rare, have been reported in yeasts such as the *Saccharomyces* hybrid VIN7 (Borneman et al. 2012), *Brettanomyces* (Borneman et al. 2014) and *S. pastorianus* group I/Saaz-type strains, which are believed to derive from multiple independent ancestral hybridisation events (Langdon et al. 2019). Phylogenetic analysis was performed to investigate whether the *H. opuntiae* x *H. pseudoguilliermondii* hybrids observed to date share a single ancestral hybridisation event.

Phylogenetic reconstruction of the *H. opuntiae* haplotypes, including the *H. opuntiae* isolates sequenced in this study, revealed evidence for at least three independent hybridisation events. Notably, the *H. opuntiae* haplotypes from two Australian hybrids were more closely related to an African isolate than Australian *H. opuntiae* isolate sequenced in this study (Figure 2b). These findings confirm that hybridisation between these species occurred independently and likely serves as an adaptive strategy to novel environments, a phenomenon observed in hybrids of other yeast genera (Gallone et al. 2019).

Ploidy estimations of the remaining sequenced isolates indicated that most strains were diploid, with only three *H. uvarum* isolates and one *H. guilliermondii* isolate exhibiting observable chromosomal aneuploidies (Table S4). Aneuploidies in yeast have been shown to arise in response to biotic or abiotic stress and, like the formation of interspecific hybrids, are often associated with adaptation to industrial fermentative environments such as those encountered in the production of beer and wine (Gilchrist and Stelkens 2019, Linder et al. 2017).

A recent study on the genetic diversity of *S. cerevisiae* isolates from spontaneous wine fermentations showed a high frequency of chromosomal instability, with 79% of isolates displaying chromosomal aneuploidies (Ward et al. 2024). Although both *Hanseniaspora* and *S. cerevisiae* occupy similar ecological niches, a comparison of aneuploidy rates suggests that *Hanseniaspora* spp. possess a relatively stable genome compared to *S. cerevisiae*. High genome stability is particularly surprising considering that *Hanseniaspora* species in general, but especially those in the FEL clade which includes *H. uvarum*, have lost a substantial cohort of genes predicted to be involved in DNA damage checkpoints and spindle checkpoint regulation, processes typically associated with genomic instability (Castro et al. 2005, Costanzo et al. 2004, Huang et al. 2000). These findings suggest that despite extensive losses of genes related to cell-cycle and DNA repair processes, *Hanseniaspora* have evolved alternative mechanisms to maintain chromosomal stability. As previously proposed, this stability may be linked to compensatory gene losses or other adaptations (Steenwyk et al. 2019). Additionally, chromosomal aneuploidies may not confer a sufficient selective advantage to become fixed within *Hanseniaspora* populations.

There was no evidence of heterozygosity or shifts in read depth in the *H. guilliermondii* isolates, suggesting that they were either haploid or homozygous diploids. A previous investigation of the genetic diversity of *Hanseniaspora* species using MLST markers also reported complete homozygosity in isolates of *H. guilliermondii*, which were suggested to be diploid based on flow cytometry (Saubin et al. 2020).

To investigate the underlying cause of the absence of heterozygous sites in *H. guilliermondii* isolates, the structure of the mating loci were specifically examined, as heterozygosity at this locus would still be expected in an otherwise homozygous diploid yeast background (due to autodiploidisation) (Magwene 2014). By inferring mating-type through protein similarity to MATa2 and MATalpha1 of *H. opuntiae* (Saubin et al. 2020), it was possible to infer complete homozygosity at the mating locus in all of the *H. guilliermondii* isolates sequenced in this study (Figure S5, Table S5). All but one contained the MATalpha genotype, with the single isolate that displayed the MATa genotype also having an inversion within this locus (Figure S5), a feature previously reported in an isolate of *H. guilliermondii* (Saubin et al. 2020). This combined evidence suggests that the *H. guilliermondii* isolates are likely haploid. However, the observed genetic diversity within the population, combined with the presence of both mating types in wild isolates, suggests active sexual recombination in the environment.

For the other sequenced diploid species (*H. uvarum*, *H. opuntiae*, *H. valbyensis*, *H. osmophila* and *H. vineae*), all but two of the *H. uvarum* isolates were heterozygous (MATa/MATalpha) at the mating locus (Table S5). Synteny analysis of the haplotypes from the long-read genome assemblies confirmed the presence of an inversion between the MATa and MATalpha alleles in species belonging to the FEL clade (Figure S5), including in *H. opuntiae*, which was previously thought to contain both mating types within a single locus (Ryan et al. 2024, Saubin et al. 2020). Reanalysing the short-read sequences of the previously published reference genome for *H. opuntiae* AWRI 3578 (Sternes et al. 2016) revealed a misassembly in the mating locus, which likely resulted from the collapse of both haplotypes, leading to the colocation of MATa and MATalpha alleles. In contrast, no inversions were observed in the mating loci of *H. osmophila* and *H. vineae* (Figure S5).

The presence of both mating types in all the species investigated suggests that sexual recombination is likely occurring. Although no sporulation was observed in a prior investigation of grape- and wine-associated *H. uvarum* isolates (Albertin et al. 2016), closely related species have demonstrated the ability to sporulate (Čadež et al. 2021, Ryan et al. 2024). Furthermore, the investigation into recombination, using linkage disequilibrium (LD) decay analysis, revealed similar rates of LD decay in wild *H. uvarum* populations compared to wild *S. cerevisiae* populations (Figure S6) (Ward et al. 2024), suggesting extensive recombination.

Differences in heterozygosity rates and population genetic diversity were observed between species, with *H. valbyensis* isolates displaying notably higher genome-wide heterozygosity and genetic diversity (Figure S1b and c). Although all isolates were obtained from the same environment, the elevated heterozygosity and genetic diversity in *H. valbyensis* suggest a more complex gene-flow history than other *Hanseniaspora* species.

*H. valbyensis* has been reported as a frequently occurring yeast in remote areas of Australia, suggesting this species occupies a broader range of ecological niches than other *Hanseniaspora* species (Varela et al. 2023, Varela et al. 2020). While speculative, the high genetic diversity observed in grape- and wine-associated *H. valbyensis* isolates may result from active recombination with isolates originating from diverse sources. Genome sequencing of *H. valbyensis* isolates from these remote areas, as well as isolates obtained from other continents, will enable the investigation of whether recent admixture events with divergent populations underlie the large heterozygosity and genetic diversity of these isolates.

### 3.2 Genomic synteny and orthology of the FEL and SEL lineages

*Hanseniaspora* species are notable for having some of the smallest genomes among Saccharomycotina yeasts, with species in the FEL clade possessing the smallest genomes and lowest gene numbers within the genus (Steenwyk et al. 2019). Compared to *S. cerevisiae*, FEL species have lost a larger number of genes associated with carbon metabolism and DNA repair than the SEL. However, the main genetic differences between the FEL and SEL clades remain unclear, as most studies have focused on well-described orthologues in *S. cerevisiae*. Additionally, the absence of chromosome-level genome assemblies for *Hanseniaspora* species has hindered the investigation of ancient chromosomal rearrangements that might explain the notably smaller genomes observed in the FEL clade.

To identify genetic differences between the FEL and SEL lineages, chromosome-level genome assemblies were generated for six *Hanseniaspora* species (Figure 3a). Gene collinearity, as well as shared and unique gene families across these species were then estimated using an orthology-based approach. Gene-based syntenic analysis revealed high chromosomal conservation within the two major phylogenetic clades (Figure 3a), with all species within each clade displaying seven highly syntenic chromosome-level contigs. However, synteny analysis between species in the FEL and SEL clades identified extensive chromosomal rearrangements, including three major translocation events involving chromosomes 3, 5, 6, and 7 (Figure 3a). These rearrangements were conserved across all FEL species, suggesting an ancestral origin.

**Figure 3.**
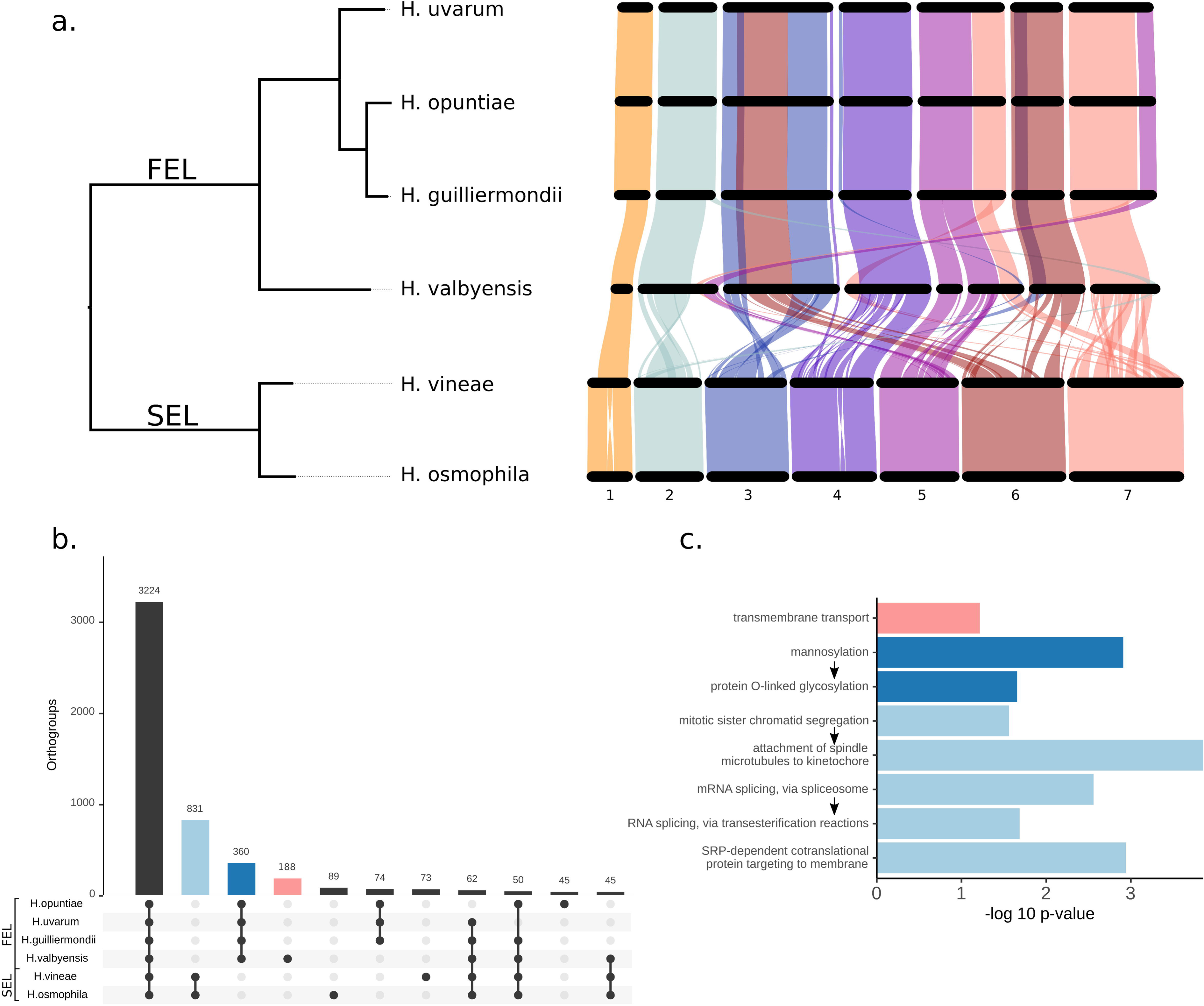
Orthology inference between *Hanseniaspora* species. **a.** Gene-based syntenic mapping between the long-read genome assemblies of six *Hanseniaspora* species. Ribbon colouring is on the basis of *H. osmophila* chromosome locations. **b.** The distribution of shared and unique orthogroups between *Hanseniaspora* species. Singleton genes not forming part of an orthogroup are also included in the total counts. Only clusters containing a minimum of 40 orthogroups are shown. **c.** Significantly enriched GO biological process categories within orthogroups. Bar colouring indicates orthogroups used for the enrichment analysis based on panel b. Arrows indicate parent-child GO terms.

Reciprocal chromosomal rearrangements have been identified as a major mechanism of reproductive isolation in yeast (Fischer et al. 2000, Hou et al. 2014), indicating that these translocations could have contributed to the evolutionary split between the SEL and FEL clades. Despite these differences, the FEL and SEL clades, which are estimated to have diverged approximately 95 million years ago (Steenwyk et al. 2019), retain a relatively high level of overall chromosomal synteny.

Additionally, investigation into the chromosomal location of orthogroups unique to species within the SEL clade, uncovered no evidence of association with the number of translocations and the number of lost genes in the FEL. This finding suggests that the extensive gene loss observed in the FEL clade is not directly linked to the specific chromosomal rearrangements that delineate the two clades.

Orthology inference between species identified 3,224 shared orthogroups across all species, with 832 and 360 orthogroups unique to the SEL and FEL, respectively (Figure 3b). *H. valbyensis* contained an unusually high number of unique orthogroups compared to other *Hanseniaspora* species (Figure 3b). To assess the biological relevance of these orthogroups, we performed GO enrichment analysis for each cluster of orthogroups (Figure 3c). Orthogroups unique to *H. valbyensis* were significantly enriched in genes involved in transmembrane transport, including genes with predicted functions as amino acid and multidrug transporters (Table S6). These genes may provide *H. valbyensis* with the ability to occupy a broader range of ecological niches than other *Hanseniaspora* species and could be linked to the higher genetic diversity and heterozygosity observed specifically in this species (Figure S1b and c).

Significantly enriched GO biological processes were also observed in orthogroups unique to the SEL clade, including those involved in chromatid segregation, mRNA splicing, and signal recognition particle (SRP)-dependent co-translational protein targeting (Figure 3c, Table S7). A recent study identified mRNA splicing and chromosome segregation as conserved gene loss events in fast-evolving lineages across three orders of the Saccharomycotina (Feng et al. 2024), suggesting that loss of these processes may represent a conserved mechanism of adaptation in yeast. Among the mRNA splicing genes, a total of 27 unique orthogroups were observed in the SEL that were associated with the pre-mRNA splicing pathway, which is primarily involved in intron removal and exon joining to form mature mRNA (Table S7) (Wahl et al. 2009). Despite this, species within both the FEL and SEL contain a comparable proportion of genes predicted to have introns (*H. uvarum*: 7.7% of genes contain at least one intron, *H. opuntiae*: 14.5%, *H.guilliermondii*: 3.3%, *H. valbyensis*: 10.9%, *H. vineae*: 12.4% and *H. osmophila*: 9.3%). Furthermore, in *H. uvarum*, a member of the FEL, 313 genes were predicted to contain introns, and these were highly enriched in genes encoding important structural constituents of the ribosome (GO:0003735, adj. pvalue 1.2×10^-9^), as also reported in *S. cerevisiae* (Parenteau et al. 2011). This enrichment suggests an active splicing capability. The apparent maintenance of this capability, despite widespread gene loss, indicates that *Hanseniaspora* species within the FEL may possess an atypical spliceosome, warranting further investigation.

Orthogroups unique to the SEL clade were also enriched in genes involved in the spindle checkpoint mechanism and chromosome segregation (Figure 3c). Notably, *H. uvarum* has demonstrated unusual cell cycle progression (Haase et al. 2024) and a faster growth rate than *S. cerevisiae* (Haase et al. 2024, Onetto et al. 2025). This accelerated growth may be linked to the absence of essential cell-cycle checkpoint mechanisms, potentially providing *H. uvarum* with a competitive advantage in fermentative environments (Onetto et al. 2025). Furthermore, orthogroups involved in the SRP pathway, which targets proteins to the secretory pathway, were absent in the FEL lineage, suggesting that species in the FEL may utilise an alternative secretory strategy, as observed in *S. cerevisiae* after loss of the SRP pathway (Mutka and Walter 2001).

### 3.3 The pangenome of *H. uvarum* is characterised by copy number variants

*H. uvarum* is considered the most abundant member of the genus on grape surfaces and in the early stages of grape wine fermentation (Liu and Howell 2021, Onetto et al. 2024). It is also commonly isolated from the surface of several fruit species other than grape (van Wyk et al. 2024). Consequently, a larger number of isolates initially identified as *H. uvarum* through ITS sequencing were sequenced in order to investigate gene content variation in this agriculturally and oenologically relevant species.

A pangenome-graph approach was employed to estimate the extent of gene content variability within the population of *H. uvarum* isolates. A reference-guided pangenome graph was constructed using three long-read, chromosome-scale assemblies of genetically divergent strains as references (wild-6, wild-104 and wild-115, Figure S7). Short-read sequencing data from 83 strains were then mapped to this pangenome, with gene presence-absence variation assessed by analysing read coverage across all annotated open reading frames (ORFs) in the pangenome graph.

Extensive genetic conservation was observed across the *H. uvarum* population, with only nine genes exhibiting strain-specific presence or absence (Table S8). Of these, two were located within the mating loci and corresponded to two strains homozygous for the mating-type locus (see Section 3.1).

Four genes orthologous to *ATM1*, *ADE4*, *GAL2*, and *RPL18A* in *S. cerevisiae* were identified as missing. However, these genes were duplicated in at least one reference genome, suggesting that while these strains lack the duplication, they retain another copy of each gene. Among the remaining six genes, a putative quinone-oxidoreductase homologous to *S. cerevisiae* YCR102C was identified, which was absent in 29 strains. In *S. cerevisiae*, YCR102C has been linked to increased ethanol production and resistance to acid stress (Chen et al. 2019), as well as enhanced copper stress tolerance in ycr102c deletion mutants (van Bakel et al. 2005). These phenotypes may provide an adaptative response to the wine environment. Furthermore, two additional genes, homologous to *BIT2* (a non-essential subunit of TORC2, a regulator of plasma membrane homeostasis) and *CAM1* (a non-essential calcium phospholipid-binding transcriptional coactivator of MXR1), were absent in two independent strains.

Given the presence-absence variations in genes identified as duplicated in reference strains, fine-scale copy-number variations (CNV) were further studied across the sequenced isolates using mean-read shifts in read-depth data (Abyzov et al. 2011). Analysis of gene-specific CNVs relative to the reference strain (wild-6) revealed substantial variation between strains, with isolates wild-167, wild-15, and wild-144 exhibiting the highest CNV levels (Figure 4a). These differences were largely attributed to chromosome aneuploidies affecting chromosomes 1, 3 and 5 (Figure 4b). Among the remaining strains, the number of genes with CNVs ranged from 11 (wild-104) to just one, with gene duplications outnumbering deletions overall (Figure 4a).

**Figure 4.**
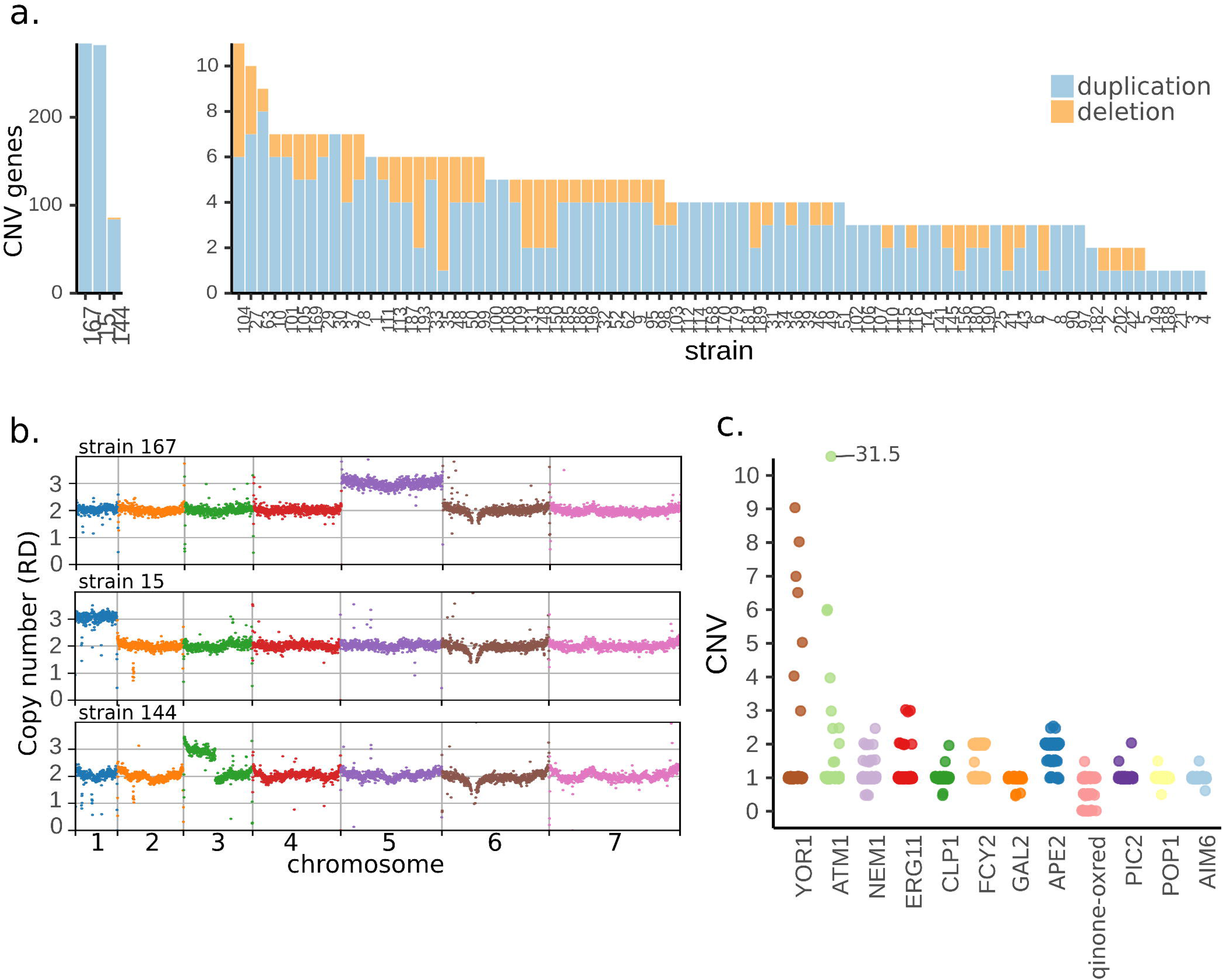
Pangenome of *H. uvarum*. **a.** Number of gene copy number variants (CNV) identified in the sequenced *H. uvarum* isolates. **b.** Read depth showing aneuploidies identified in three H. uvarum isolates. **c.** Specific gene copy number variations (2N based, CNV of 1 = 2 copies) identified in the sequenced *H. uvarum* isolates. Only genes showing CNV in at least two strains and with predicted function are presented. Gene IDs are based on homology to *S. cerevisiae* s288c. Details on all the identified CNVs are available in Table S9.

Using the mean-read shift approach, copy numbers were estimated for those genes exhibiting CNV (Figure 4c). Genes orthologous to *S. cerevisiae YOR1* and *ATM1* displayed the most pronounced CNV differences, with one strain carrying up to 31.5 copies (63 total copies) of *ATM1* (Figure 4c). *YOR1* encodes an ATPase-coupled xenobiotic transmembrane transporter, initially identified for its role in oligomycin resistance in *S. cerevisiae* (Katzmann et al. 1995). Later studies linked this transporter to sensitivity to various fungicides, herbicides, pesticides, antibiotics, and antiseptics (Rogers et al. 2001). Given that *H. uvarum* is consistently found in high abundance on fruit surfaces (van Wyk et al. 2024), particularly on grapes (Liu and Howell 2021, Onetto et al. 2024), this CNV may reflect an adaptive response to continuous exposure to agricultural xenobiotics. Similar CNV-driven adaptations have been observed in response to other environmental stresses in yeast, such as SO_2_ and Cu (Bartel et al. 2021, Steenwyk and Rokas 2018).

*ATM1* encodes a highly conserved and well characterised ABC transporter located in the mitochondrial inner membrane, essential for the assembly of cytosolic-nuclear iron/sulfur (Fe/S) clusters, critical cofactors in various metabolic pathways (Kispal et al. 1999, Leighton and Schatz 1995, Srinivasan et al. 2014). Given the fundamental role of Atm1 in cellular function, the underlying cause of its extensive CNV among strains remains unclear. Despite this, CNVs and overexpression of *ATM1* along with *ERG11*, which also exhibited CNVs in *H. uvarum* isolates (Figure 4c), have been directly linked to azole antifungal resistance in *Malassezia restricta*, *Candida albicans*, and *Cryptococcus neoformans* (Cowen et al. 2000, Selmecki et al. 2008). The links to antifungal resistance in other yeast species suggests that increased *ATM1* copy number in *H. uvarum* may contribute to an unidentified adaptive advantage, potentially influencing a resistance mechanism.

Among the remaining genes, several play key roles in nutrient transport and metabolism, possibly reflecting adaptation to the nutritional stresses associated with the grape and wine environment. These include *FCY2* (encoding a purine-cytosine permease), *APE2* (encoding a leucine aminopeptidase, 2 copies in reference), *GAL2* (encoding a galactose permease, 4 copies in reference) and *PIC2* (encoding a mitochondrial copper and phosphate carrier). Similar gene amplifications linked to nutrient acquisition and metal homeostasis have been observed in *S. cerevisiae* (Steenwyk and Rokas 2018), suggesting that CNVs in *H. uvarum* may contribute to phenotypic diversity and enhanced tolerance to common wine fermentation stresses (see later section).

Further investigations are required to elucidate the functional consequences of these CNVs, particularly their potential role in stress resistance, metabolic adaptation, and ecological success in fermentation environments.

### 3.4 The selection landscape of *Hanseniaspora* species

Recent investigations into the evolutionary history of *Hanseniaspora* species estimate that the FEL and SEL lineages diverged approximately 95 million years ago (Steenwyk et al. 2019). While the selective pressures driving this divergence remain unknown, the emergence of this genus coincides with the Early Cretaceous radiation of flowering and fruiting angiosperms (Crane et al. 1995, Zuntini et al. 2024), including grapes (Wikström et al. 2001). *Hanseniaspora* species of the FEL lineage are commonly isolated from fruit (Chanprasartsuk et al. 2010, Prada and Pagnocca 1997, Ramirez-Castrillon et al. 2019, Vadkertiová et al. 2012, Vegas et al. 2020) and extracted juices (Chanprasartsuk et al. 2010, Lorenzini et al. 2018, Onetto et al. 2024, Satora and Tuszynski 2005), while SEL species are infrequently observed (Liu and Howell 2021, Onetto et al. 2024, Prada and Pagnocca 1997). Experimental evidence also demonstrates that FEL *Hanseniaspora* have a higher abundance than SEL species during fermentation (Liu and Howell 2021, Onetto et al. 2024, Snyder et al. 2024). Therefore, the concordance between the timing of FEL/SEL divergence and the diversification of fruiting plant species, along with the abundance and success of FEL species within the fruit/juice microbiome, suggests that bifurcation of FEL and SEL may have occurred as a result of colonisation of the fruit niche by early FEL species.

The colonisation of a new environment imposes selection pressure on a range of diverse pathways depending on the metabolites and stressors present within the new environment (Rosa et al. 2018, Rundle and Nosil 2005). To understand the selective landscape immediately after the bifurcation of FEL/SEL and before the diversification of extant FEL species, single copy orthologs were tested for episodic diversifying selection (n = 2036). It was hypothesised that genes under selection within the FEL branch may have been driven by early adaptation to the fruit niche. Genes were investigated under three hypotheses (Figure 5A): i) evidence for diversifying selection only along the FEL branch, ii) evidence for diversifying selection only along the SEL branch and iii) evidence for diversifying selection along both FEL and SEL but under different selective regimes. This identified 229 genes under diversifying selection specific to FEL (hypothesis i) and 93 with selection along both FEL and SEL branches with different selective regimes (hypothesis iii) (Table S10).

**Figure 5.**
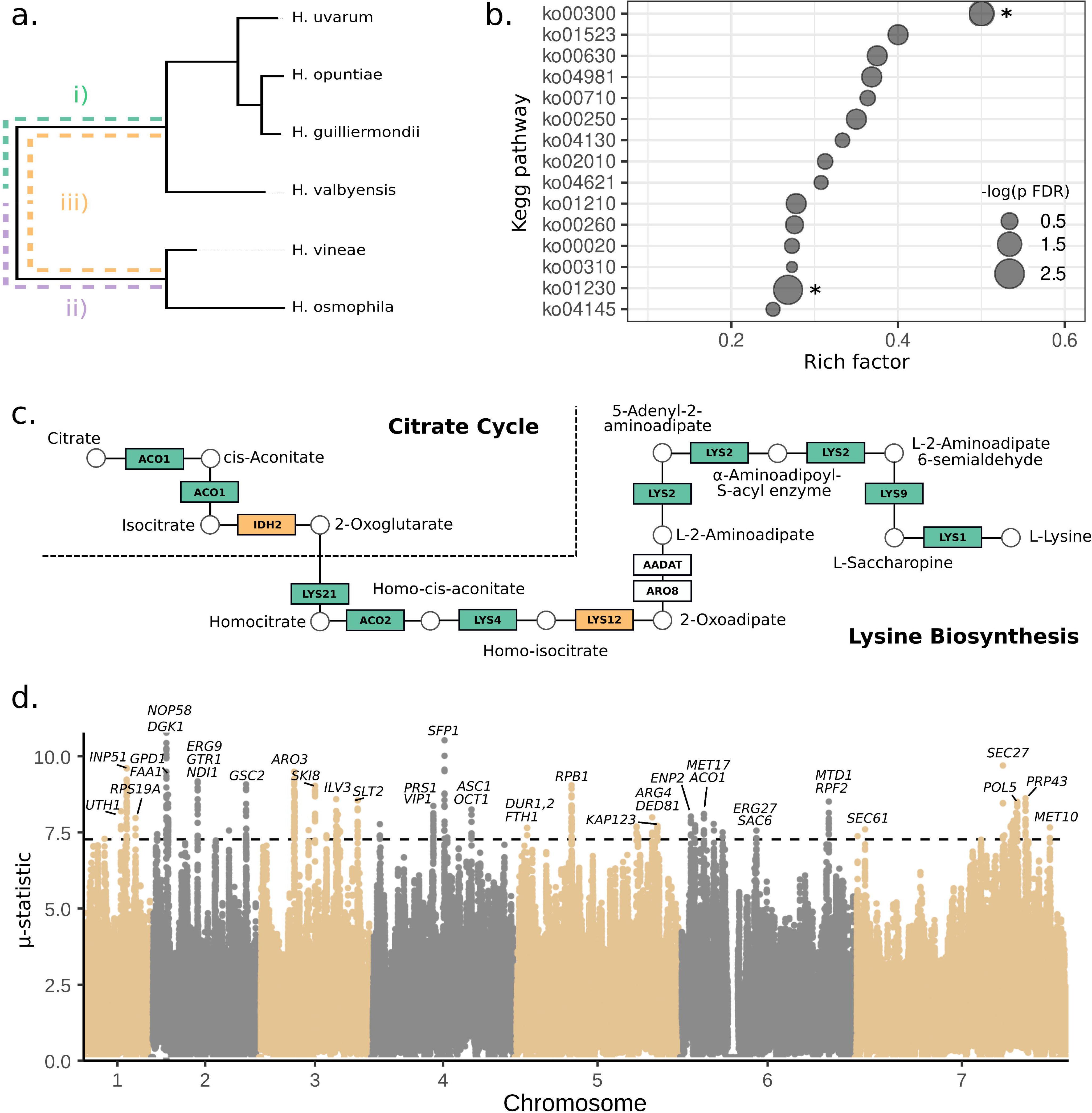
Selection signatures of *Hanseniaspora*. **a.** Selection hypotheses tested against the *Hanseniaspora* species tree generated in this study, i) episodic diversifying selection along the FEL branch, ii) episodic diversifying selection along the SEL branch, iii) episodic diversifying selection along both the FEL and SEL branch but with significantly different selective regimes. **b.** KEGG pathway enrichment with significant evidence of episodic diversifying selection along the FEL branch (hypotheses i and iii). Rich Factor represents the number of genes under selection within a given pathway and the total number of genes within that pathway in the *H. uvarum* genome. Significantly (FDR corrected p < 0.05) enriched pathways lysine biosynthesis (ko00300) and biosynthesis of amino acids (ko01230) are denoted with asterisks. **c.** KEGG enzymes catalysing the reactions from citrate to L-lysine within the citrate cycle (ko00020) and lysine Biosynthesis (ko00300) pathways. Genes under significant episodic diversifying selection are coloured depending on which hypothesis (outlined in **a.**) was significant. **d.** Distribution of the μ statistic across the genome of *H. uvarum*. ORFs overlapping with top scored regions are labelled for each chromosome. Dotted line represents the 99.9% μ statistic cutoff threshold used to select sweep regions.

Diversifying selection was identified in 165 KEGG pathways along the FEL branch (Table S10). Two KEGG pathways, lysine biosynthesis (ko00300) and biosynthesis of amino acids (ko01230), were significantly enriched when compared to the total ortholog space (p FDR < 0.05) (Figure 5B). Manual investigation of pathway maps adjacent to ko00300 revealed that nine adjacent enzymes across the citrate cycle (*ACO1* and *IDH2*) and lysine biosynthesis (*LYS1*, *LYS2*, *LYS4*, *LYS9*, *LYS12*, *LYS21* and *ACO2*) KEGG pathways were under diversifying selection (Figure 5C). This suggests that the FEL clade underwent widespread diversifying selection across almost all enzymes in the pathway from citrate to L-lysine before the diversification of the FEL clade into the extant species utilised in this study. Although it was not determined if diversifying selection across the lysine biosynthesis pathway has directly resulted in increased L-lysine production, selection may have occurred as an adaptation to a low lysine environment, such as fruit surfaces and juice (Gutiérrez-Gamboa et al. 2018, Zeng et al. 2015). The lysine biosynthesis pathway has also been shown to be involved in the yeast oxidative stress response (O’Doherty et al. 2014), converting L-lysine to polyamines to be utilised in the glutathione biosynthesis pathway (Olin-Sandoval et al. 2019). Therefore, selection in the lysine biosynthesis pathway may be an indirect adaptation to oxidative stress present within the fruit fermentative environment. Further research is needed to elucidate the physiological implications of these adaptive signals within the lysine biosynthesis pathway of FEL and to determine whether they result in metabolic differences in the utilisation of this amino acid.

*H. uvarum* has successfully dominated the grape fermentative environment being consistently observed as the most dominant non-*Saccharomyces* species during fermentation. While part of this success might be attributed to ancestral gene loss events (Haase et al. 2024) and a rapid growth rate (Onetto et al. 2025), identifying *H. uvarum* genes under recent positive selection might provide some hints as to the selective pressures that have shaped *H. uvarum*’s adaptation to the grape and wine environment. To investigate selection associated with adaptation to the wine fermentation environment, the *H. uvarum* genome was scanned for signatures of selective sweeps across a population of 84 *H. uvarum* grape/wine-associated isolates. A percentile threshold of 99.9% was applied to the μ statistic, and top scored overlapping windows were merged to detect sweep regions (Figure 5d).

A total of 78 genes were detected within these regions (Table S11), spanning multiple metabolic pathways, including carbohydrate, amino acid, and lipid metabolism, as well as genetic information processing (Table S11).

Polymorphisms in genes linked to key stress responses in grape juice and wine fermentation were under selection. For example, *MET10* and *MET17* were located within selective sweep regions (Figure 5d). These genes encode sequential enzymes in the sulphur assimilation pathway. *MET10* encodes a sulphite reductase, a key enzyme in the assimilation and detoxification of intracellular SO₂ (Nadai et al. 2016), which is the most widely used antiseptic and antioxidant in winemaking (Ribéreau-Gayon et al. 2006). Interestingly, FEL species lack orthologs of *SSU1*, the efflux pump responsible for sulphite tolerance in *S. cerevisiae* (Park and Bakalinsky 2000), suggesting that alternative mechanisms may contribute to SO₂ tolerance in *H. uvarum*.

Genes involved in sterol biosynthesis, including *ERG9* and *ERG27*, were also under positive selection (Table S11). Sterols are essential components of the plasma membrane and play a direct role in ethanol tolerance during wine fermentation (Girardi Piva et al. 2022). Additionally, *GPD1*, which encodes glycerol-3-phosphate dehydrogenase, a key enzyme in glycerol synthesis, was also under positive selection. Glycerol production is essential for yeast survival under osmotic stress, such as the high sugar concentrations found in grape must (Albertyn et al. 1994, Jiménez-Martí et al. 2011).

Overall, these findings suggest that *H. uvarum* has undergone recent genetic adaptations to withstand the challenging conditions of grape must and wine fermentation, including exposure to SO₂, ethanol, and osmotic stress. Further studies are needed to elucidate the precise molecular mechanisms underpinning these adaptations and their impact on yeast survival during grape must fermentation.

### 3.5 Phenotypic diversity of *Hanseniaspora*

Several studies have investigated how the metabolism of *Hanseniaspora* species withstand common wine-related stresses and influence the physicochemical characteristics of wine (van Wyk et al. 2024). Despite this, limited information is available on strain variability, with only a single study exploring the phenotypic diversity of a large set of *Hanseniaspora* species (Albertin et al. 2016). Given the extensive genetic diversity and copy number variations (CNVs) observed among the isolates sequenced in this study, a high-throughput methodology was employed to screen a selection of 113 isolates for a set of wine-related traits to determine whether the observed genetic diversity translates into phenotypic variability (Figure 6).

**Figure 6.**
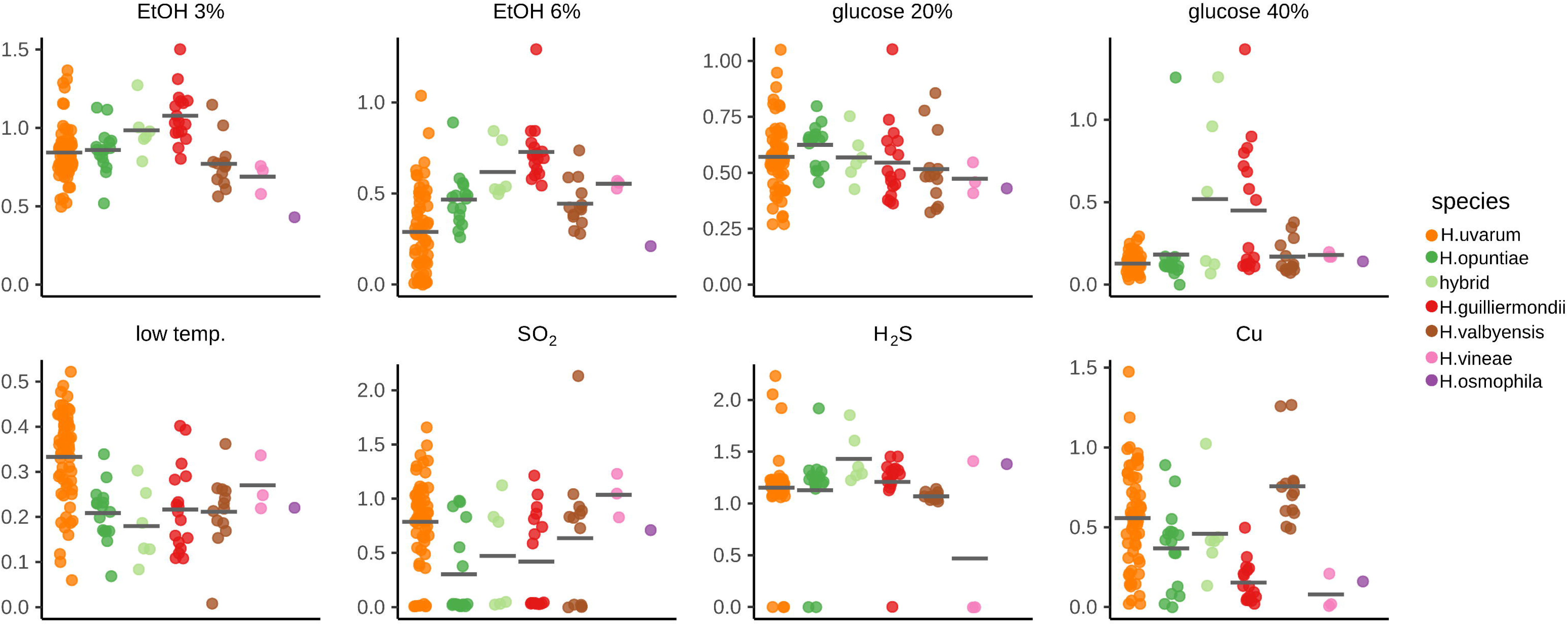
Phenotypic characterisation of *Hanseniaspora* species. Each point represents the ratio between the colony size of the phenotype tested and the control condition for each isolate. Each points represents the mean of two replicates. Grey line shows the mean for each species.

Substantial phenotypic diversity was observed across the isolates, with intraspecific variation often exceeding interspecific differences, particularly in *H. uvarum*, which had the largest number of isolates tested (Figure 6). Nevertheless, specific trends emerged for specific traits. For example, *H. guilliermondii* and the *H. opuntiae* × *H. pseudoguilliermondii* hybrid isolates had a significantly greater (adj. p-value < 0.05) resistance to ethanol and osmotic stress (40% glucose) than most other species (Figure 6, Table S12). High ethanol tolerance in *H. guilliermondii* has been previously reported, with isolates demonstrating greater ethanol resistance than *H. uvarum* and reaching tolerance levels comparable to *S. cerevisiae* (Pina et al. 2005, Pina et al. 2004). Similarly, *H. uvarum* isolates displayed significantly higher resistance to low temperatures than other species, which may reflect their ability to endure prolonged cold maceration during winemaking (Hall et al. 2017, Johnson et al. 2020). While Albertin et al. (2016) previously reported growth of most *H. uvarum* isolates at 12°C, variability among isolates was not addressed.

Two traits, tolerance to copper and SO₂, produced unexpected results. In *S. cerevisiae*, these traits are associated with specific genes, including the metallothionein protein *CUP1* and the sulfite efflux pump *SSU1* (Fogel et al. 1983, Park and Bakalinsky 2000). While a *CUP1* homolog is absent in all *Hanseniaspora* species examined, significant differences in copper tolerance were observed, with *H. valbyensis* exhibiting the highest mean resistance (Figure 6), suggesting alternative mechanisms for copper tolerance in this genus. Homologs of *SSU1* are present in SEL species, including *H. vineae* and *H. osmophila*, consistent with the higher mean SO₂ tolerance observed in *H. vineae*. However, SO₂ tolerance varied widely across species, indicating that alternative mechanisms could also contribute to SO₂ resistance in *Hanseniaspora*.

Considering the unexpectedly large phenotypic differences among *H. uvarum* isolates and the availability of whole-genome sequencing data, potential genotype-phenotype associations were investigated using a GWAS approach. Signals were assessed for each phenotype and genomic loci were examined for variations within ORFs that might be associated with specific traits. GWAS analysis identified a potential association between copper tolerance and a synonymous variant in a homolog of *S. cerevisiae YCF1*, which encodes a vacuolar glutathione S-conjugate ABC transporter involved in the detoxification of bivalent metal cations (Gueldry et al. 2003, Li et al. 1997, Szczypka et al. 1994) (Figure S8). The mutation (997C>T) replaces the glycine codon GGC with GGT, the preferred glycine codon in *H. uvarum* (Figure S8b, c) (Zavala et al. 2024). Future studies should investigate whether this synonymous mutation and the *YCF1* homolog play a role in the copper tolerance phenotype in *Hanseniaspora*, as has been observed for synonymous mutations in yeast (Shen et al. 2022).

Overall, these findings highlight the extensive phenotypic diversity among *Hanseniaspora* isolates and understanding the mechanisms behind these strain-level differences is essential for evaluating the role of *Hanseniaspora* in fermentation dynamics and could inform future applications in winemaking, particularly in the selection of beneficial strains for modulating fermentation outcomes.

## 4. Conclusions

This study explored the genomic and phenotypic diversity of *Hanseniaspora* species isolated from grape and wine fermentation environments. Long-read genome assemblies identified ancestral, lineage-specific chromosomal translocation events that may have contributed to the evolutionary split between the SEL and FEL clades. Species in the FEL clade have undergone large-scale, conserved gene loss in pathways related to chromatid segregation, mRNA splicing, and signal recognition particle-dependent protein targeting. Despite this, *Hanseniaspora* genomes exhibit remarkable chromosomal stability, suggesting the evolution of alternative genome maintenance mechanisms. Pangenome analysis of *H. uvarum* revealed extensive CNVs, particularly in genes associated with xenobiotic resistance and nutrient transport, indicating ongoing adaptive diversification within this species. Selection analyses following the FEL/SEL divergence highlighted the lysine biosynthesis pathway as a key target of early FEL adaptation. High-throughput phenotypic screening further uncovered substantial intraspecific variation in wine stress tolerance traits, including resistance to ethanol, SO₂, copper, and low temperature. Together, these findings position *Hanseniaspora* as a powerful model for studying genome evolution, ecological adaptation, and functional innovation in yeast

## 5. Acknowledgements

This work was supported by Wine Australia, with levies from Australia’s grape growers and winemakers and matching funds from the Australian Government. The AWRI is a member of the Wine Innovation Cluster (WIC) in Adelaide.

