## Supplementary material for "Genetic and phenotypic diversity of wine-associated *Hanseniaspora* species": Figure S1, Figure S2, Figure S3, Figure S4, Figure S5, Figure S6, Figure S7, Figure S8

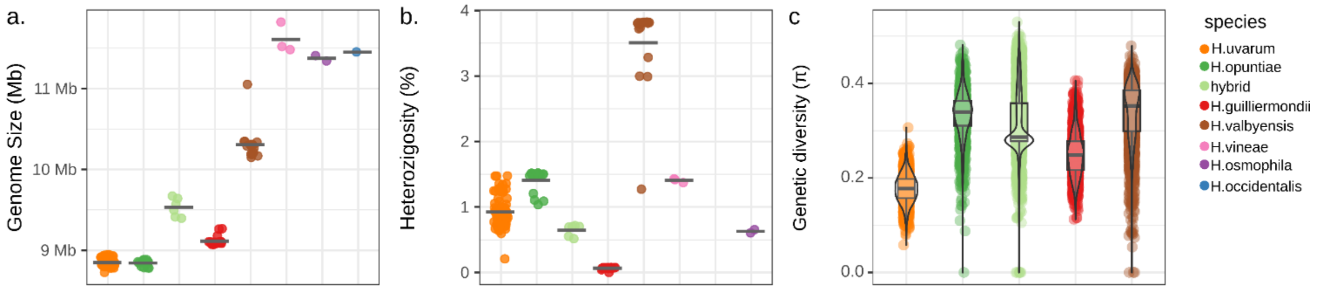


Figure S1. The genetic diversity of *Hanseniaspora* isolates. **a.** Genome size of the short-read genome assemblies after contig redundancy reduction. **b.** Estimated heterozygosity for each species. **c.** Estimated nucleotide diversity in 5 kb sliding windows.

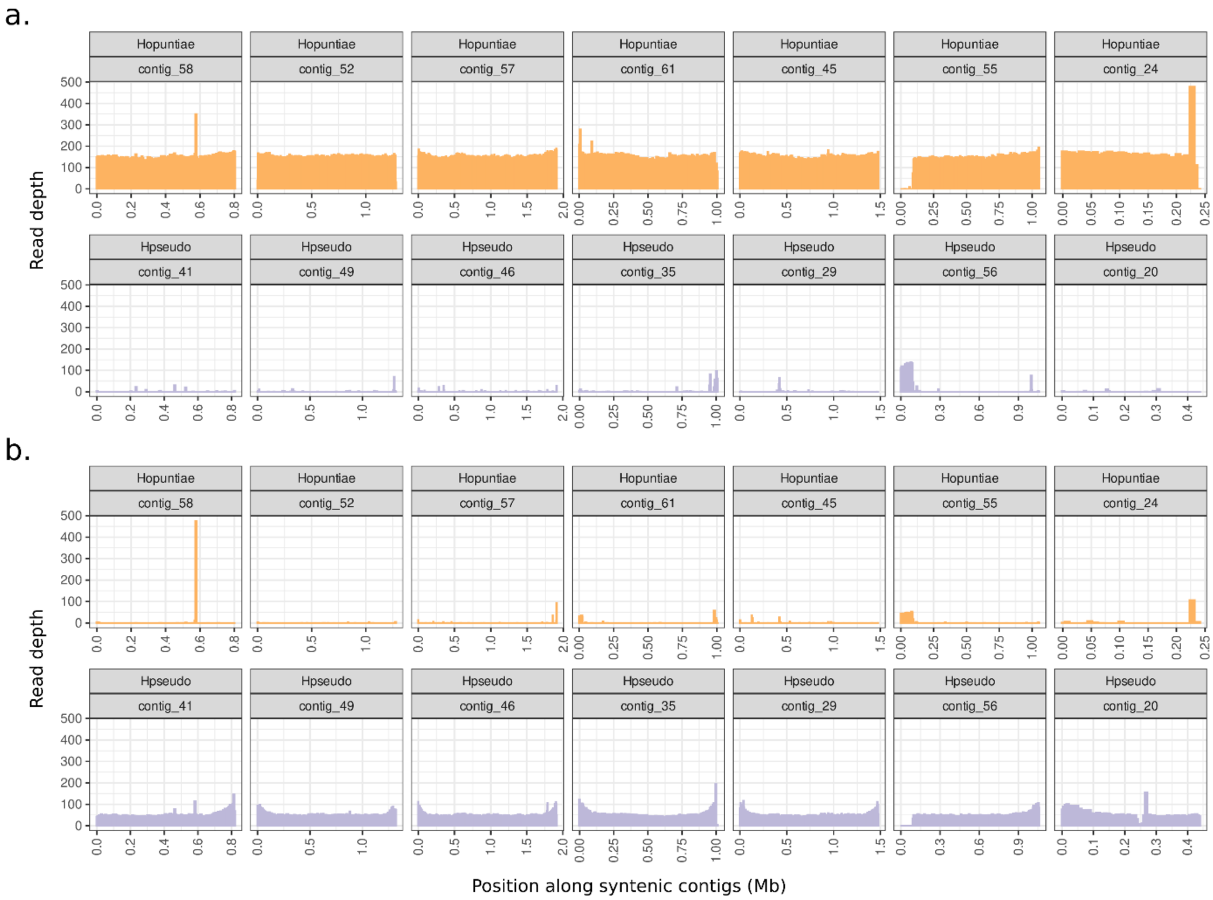


Figure S2. Raw read depth mapped to the wild-127 *H. opuntiae* x H. *pseudoguilliermondii* hybrid reference genome. Syntenic contigs of *H. opuntiae* and *H. pseudoguilliermondii* origin have been separated to show coverage of the *H. opuntiae* and *H. pseudoguilliermondii* reads. **a.** *H. opuntiae* (wild-121) coverage. 97.7 % of the reads mapped to the wild-127 *H. opuntiae* haplotype. **b.** *H. pseudoguilliermondii* (SRA: SRR16988876) coverage. 96.5 % of the reads mapped to the wild-127 *H. pseudoguilliermondii* haplotype.

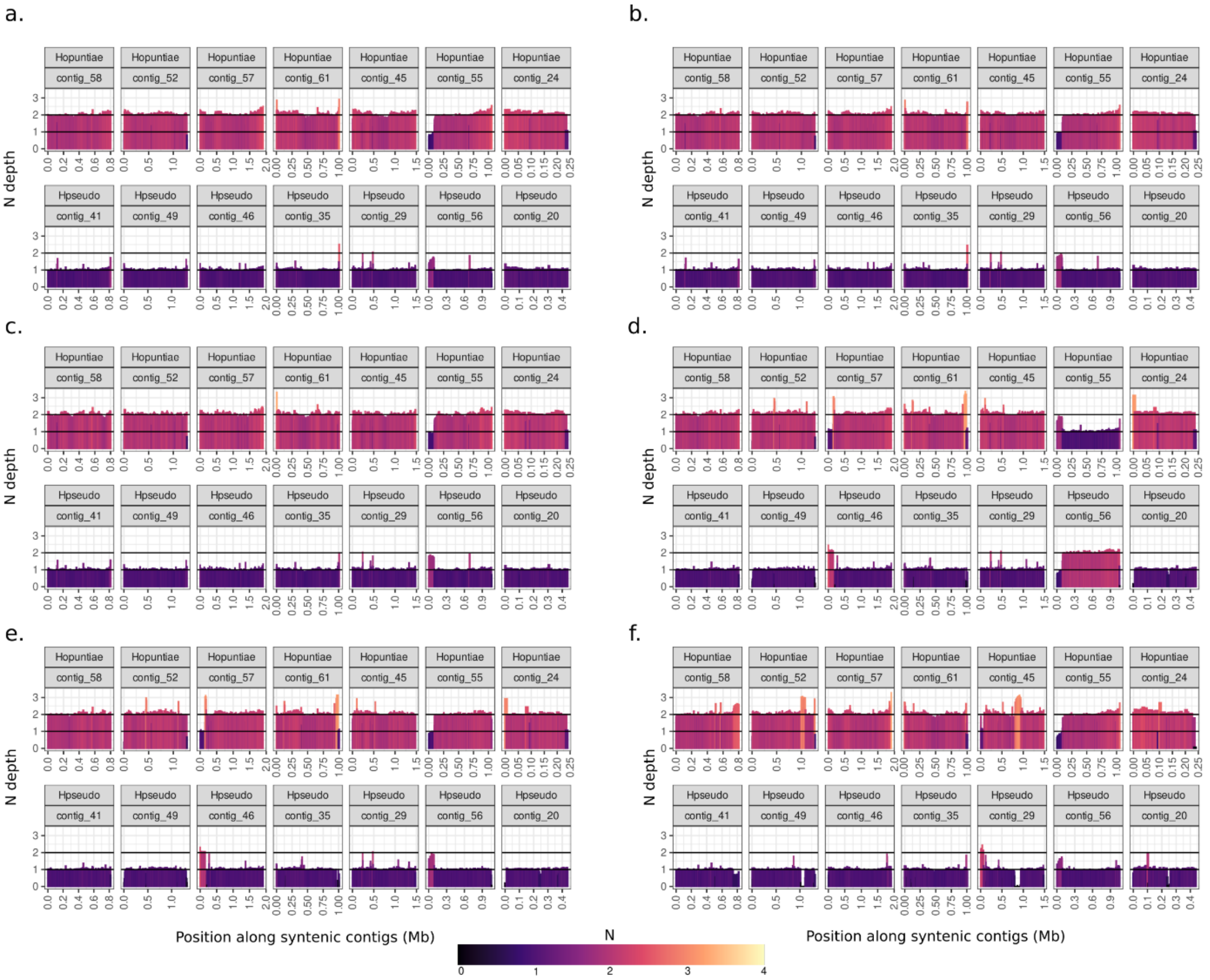


Figure S3. Relative 2N read depth of *H. opuntiae* x *H. pseudoguilliermondii* hybrids sourced from the wine environment in this study mapped to the wild-127 *H. opuntiae* x *H. pseudoguilliermondii* reference genome. Syntenic contigs of *H. opuntiae* and *H. pseudoguilliermondii* origin have been separated to show ploidy of the *H. opuntiae* and *H. pseudoguilliermondii* genome haplotypes within the hybrids. **a.** Relative coverage of wild-120. **b.** Relative coverage of wild-127. **c.** Relative coverage of wild-128. **d.** Relative coverage of wild-44. **e.** Relative coverage of wild-45. **f.** Relative coverage of wild-61.

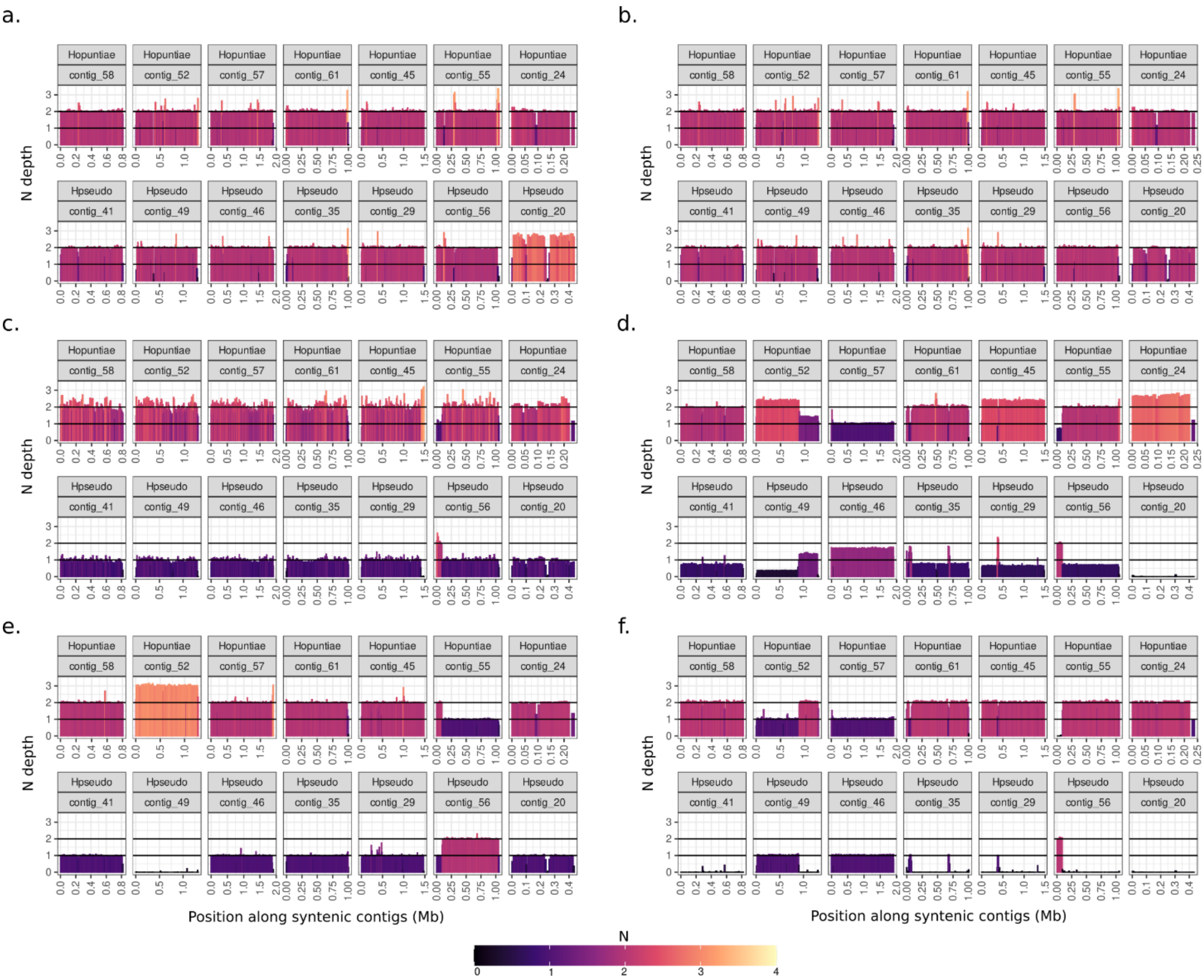


Figure S4. Relative 2N read depth of publicly available *H. opuntiae* x *H. pseudoguilliermondii* hybrids mapped to the wild-127 *H. opuntiae* x *H. pseudoguilliermondii* reference genome. Syntenic contigs of *H. opuntiae* and *H. pseudoguilliermondii* origin have been separated to show ploidy of the *H. opuntiae* and *H. pseudoguilliermondii* genome haplotypes within the hybrids. **a.** Relative coverage of CLIB_3263 (SRA: ERR3456262). **b.** Relative coverage of CLIB_3313 (SRA: ERR3456263). **c.** Relative coverage of CLIB_3101 (SRA: ERR3456264). **d.** Relative coverage of CCY46_1-3 (SRA: ERR3456265). **e.** Relative coverage of DBVPG_5828 (SRA: ERR3456266). **f.** Relative coverage of CCY46_1-3a (SRA: ERR3456268).

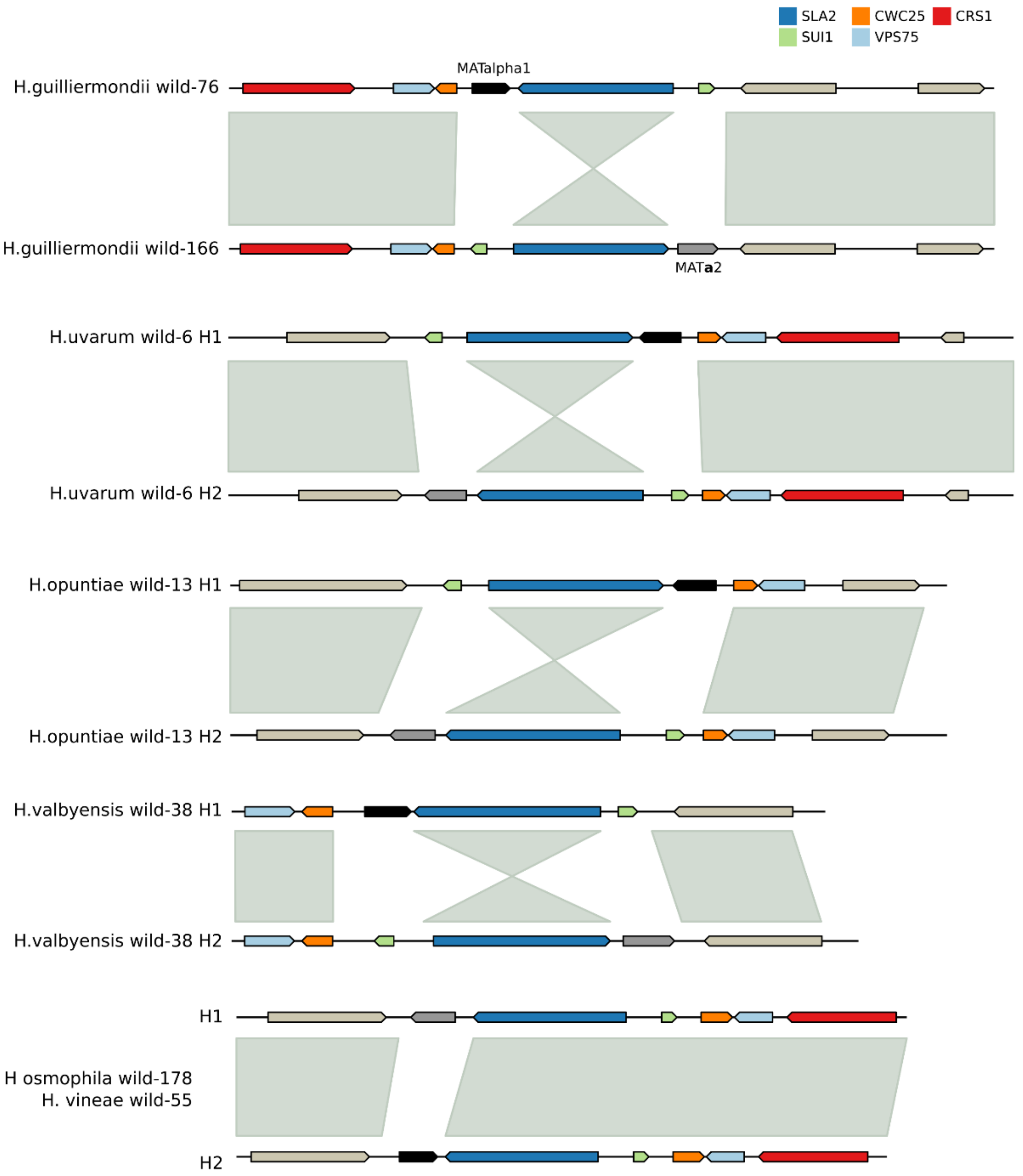


Figure S5. The predicted mating locus in the long-read genome assemblies of *Hanseniaspora* species. Syntenic alleles within each species are shown for all species except *H. guilliermondii* where haplotypes are shown for two different isolates.

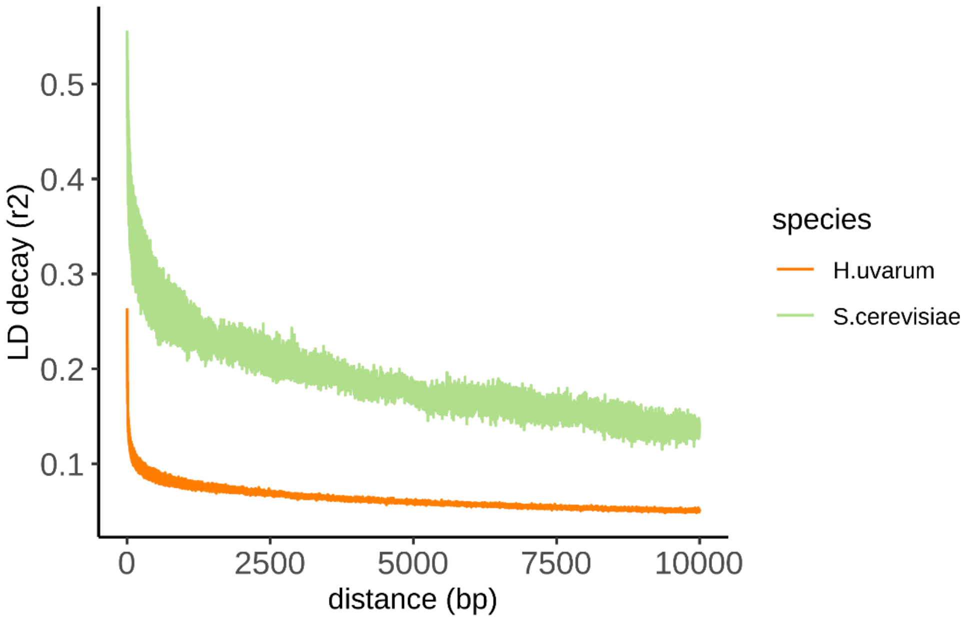


Figure S6. Decay of linkage disequilibrium (LD) for *H. uvarum* isolates and previously published spontaneous isolates of S. cerevisiae (Ward et al., 2024).

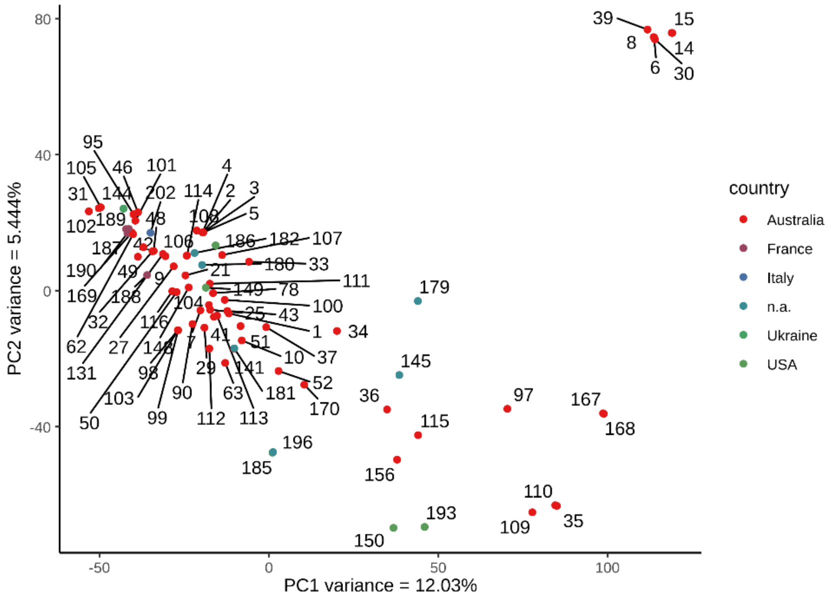


Figure S7. Principal component analysis of the SNPs identified in the *H. uvarum* isolates. Strains have been coloured based on country of isolation. n.a: not available.

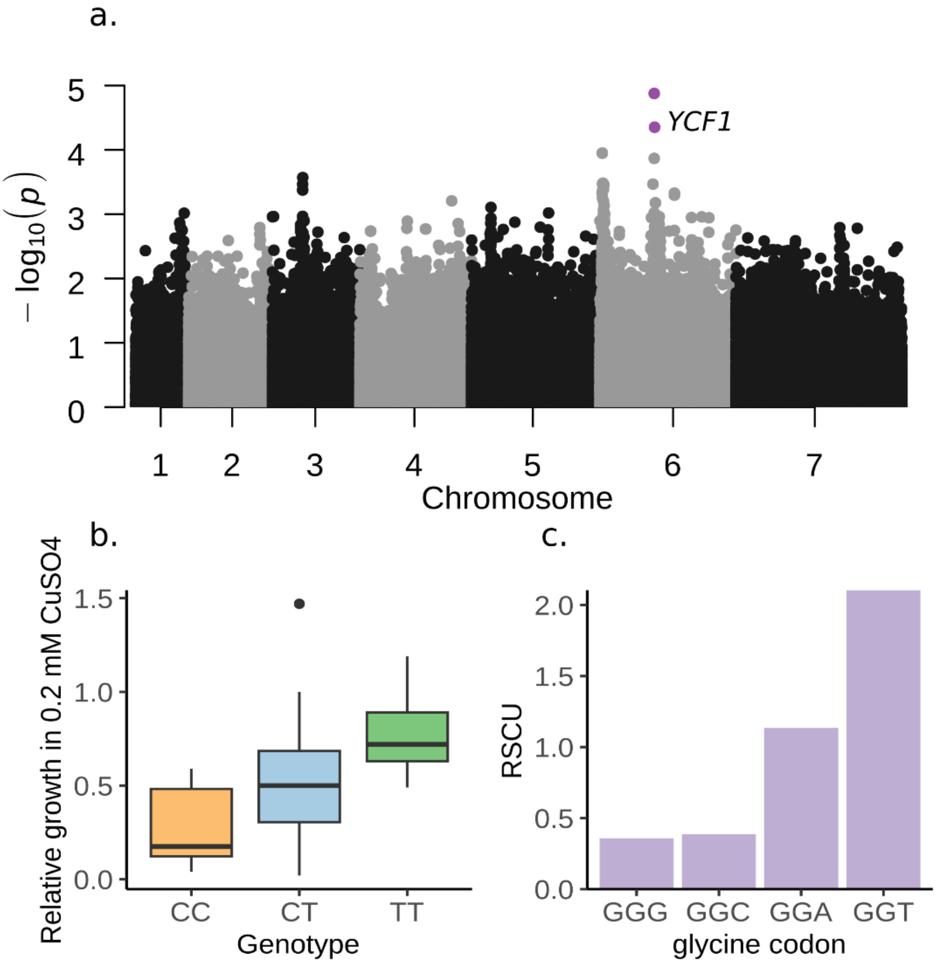


Figure S8. Genome wide association analysis of single nucleotide polymorphisms (SNPs) and the copper tolerance phenotype in *H. uvarum*. **a.** Manhattan plot of the SNPs, the x axis shows chromosomal position, and the y axis shows −log10 (P). **b.** Boxplot showing the relative growth in copper medium for the three genotypes identified in the SNP of *YCF1*. **c.** Relative synonymous codon usage (RSCU) values for glycine codons in *H. uvarum* as previously published by Zavala et al. (2024).
